# Comparing the genetic diversity of two threatened *Rhyticeros* hornbill species with contrasting geographic range sizes

**DOI:** 10.1101/2025.10.06.680625

**Authors:** N Devanand, D. K. Bharti, Pooja Yashwant Pawar, Abhishek Gopal, Sartaj Ghuman, Navendu Page, Jahnavi Joshi, Rohit Naniwadekar

## Abstract

Island endemics exhibit low genetic diversity due to founder effects and geographical isolation, increasing their vulnerability to inbreeding depression. The Narcondam Hornbill (*Rhyticeros narcondami*), restricted to the 6.8 km^2^ Narcondam island in the Andaman archipelago, has the smallest range size among hornbills, a globally threatened group of birds. We compared the genetic diversity of the Narcondam Hornbill with the Wreathed Hornbill (*Rhyticeros undulatus*) distributed from the Eastern Himalaya to Bali. We generated DNA sequences for four mitochondrial markers ; Cytochrome B (Cytb), cytochrome c oxidase subunit I (COI), NADH dehydrogenase subunit 2 (ND2) and displacement loop (D-loop) regions) from 14 Narcondam Hornbill faecal samples and for three markers (Cytb, COI and ND2) from 19 Wreathed Hornbill tissue samples from north-east India. Our results suggest markedly lower genetic diversity in the island endemic Narcondam Hornbill compared to the Wreathed Hornbill. Specifically, Cytb and COI showed no genetic variation in the island endemic, compared to three to five haplotypes in the Wreathed Hornbills. In the ND2 region, Narcondam Hornbills showed seven haplotypes among 13 samples compared to the Wreathed Hornbill’s six haplotypes among eight samples. The observed genetic diversity in the D-loop in the Narcondam Hornbill was lower than other island-endemic hornbill species in the Philippines. The extremely low genetic diversity in the Narcondam Hornbill likely stems from a small founder population and/or past hunting pressures that reduced its numbers to approximately 30% of its current population of 1000 birds, rendering the Narcondam Hornbill susceptible to environmental and human-driven changes.

## INTRODUCTION

Islands harbour evolutionarily distinct diversity, highlighting the value of conserving these habitats (Jetz et al. 2014 and Jønsson and Holt 2015). Island endemic birds are documented to have a 143 times higher extinction rate compared to continental species (Loehle and Eschenbach 2012). Large, forest-dwelling island birds are known to be particularly vulnerable to extinction (Matthews et al. 2022). Apart from threats posed by habitat loss, hunting, and invasive species, low genetic diversity can also play a key role in influencing the extinction rates of island species (Duncan and Blackburn 2007).

Dispersal events, where a small number of individuals carry only a fraction of the genetic diversity from the source population, play a key role in influencing the genetic diversity of extant populations. Such small founder populations are susceptible to a loss of genetic diversity through the process of genetic drift (Charlesworth 2009). Lower genetic diversity in island populations can further decrease due to their susceptibility to hunting, habitat loss and demographic bottlenecks due to stochastic events and geographic restriction influencing population sizes (Gimingham 1993; Frankham 1997). Low genetic diversity can threaten island populations through compromising their ability to adapt to environmental change (DeWoody et al. 2021), reducing the strength of positive selection on new advantageous alleles (Charlesworth 2009), and failure to remove deleterious alleles (Agrawal and Whitlock 2012). Mating between close relatives in a small population can also lead to inbreeding depression (Charlesworth and Willis 2009). These genetic mechanisms pose a long-term risk to island species and populations, which can ultimately lead to extinction (Fernández-Palacios et al. 2021).

Hornbills are among the largest forest-dwelling birds in the Asian tropics. Sixteen of 32 Asian hornbill species are found on islands, twelve of which are threatened (IUCN, 2025). Among island hornbills, the Narcondam Hornbill (*Rhyticeros narcondami*; weight: ∼750g) has the smallest global range as it is endemic to the 6.8 km^2^ Narcondam island in the Andaman and Nicobar archipelago, located in the Bay of Bengal, east of the Indian subcontinent. Its close relative in the Indian mainland, the Wreathed Hornbill (*Rhyticeros undulatus*; weight: ∼3kg), is migratory and has a very wide geographic range spanning from the Eastern Himalaya to Bali (Gonzalez et al. 2013) (Fig. 1). Both species are known to play a critical role in seed dispersal (Naniwadekar et al. 2019, 2021b). The two species offer contrasting examples for comparing genetic diversity. Given this background, we carried out this study with the aim of comparing the genetic diversity of the point endemic Narcondam Hornbill with the Wreathed Hornbill, which has a wide geographic range, using a combination of faecal and tissue samples.

**Figure 1.**
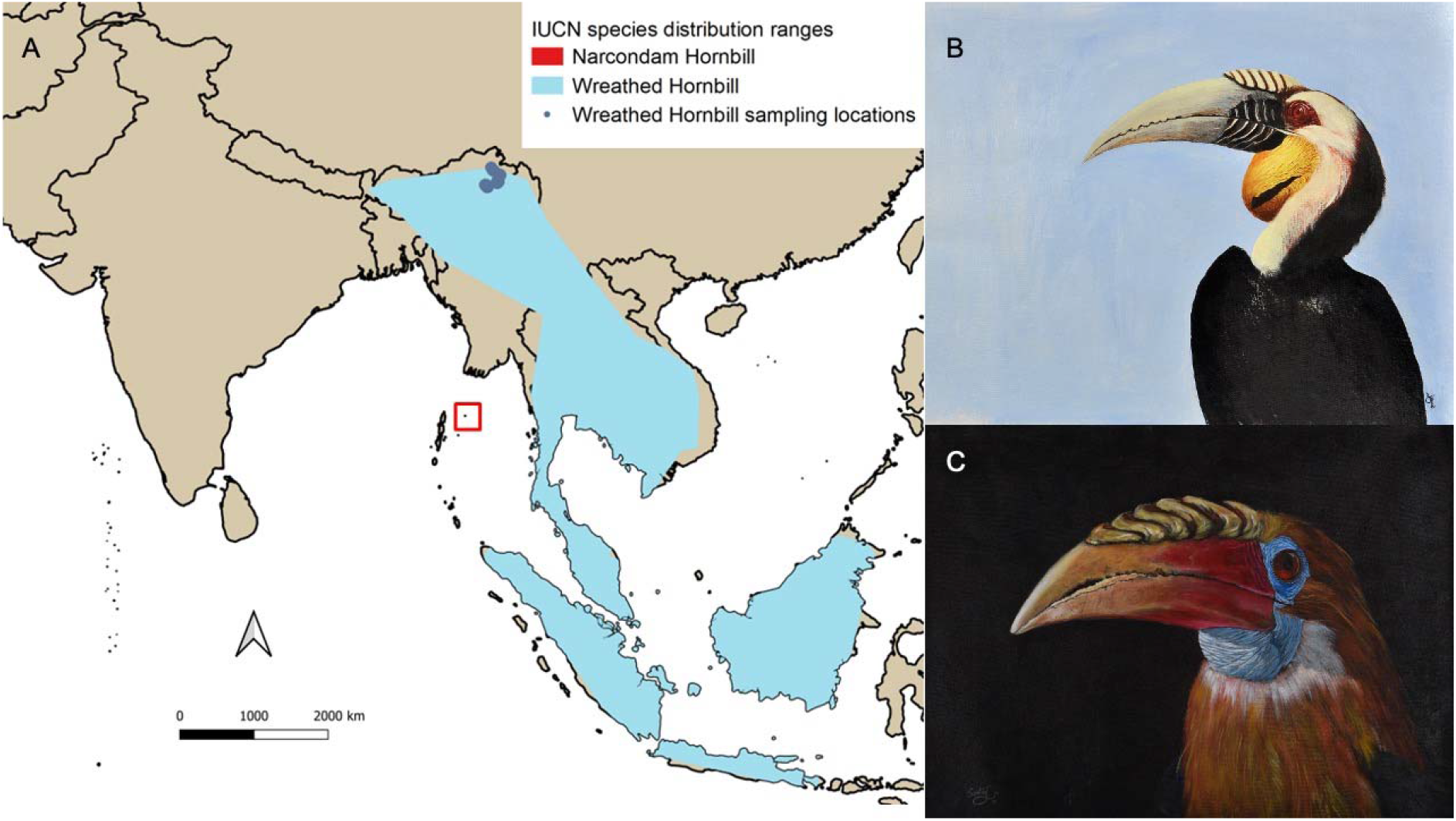
Distribution ranges and sampling sites (A) of the Wreathed Hornbill *Rhyticeros undulatus* (blue; B) and Narcondam Hornbill *Rhyticeros narcondami* (red; C). Paintings by Sartaj Ghuman.

## METHODS

### Sample Collection

RN, AG, SG and NP collected faecal samples of Narcondam Hornbills in absolute ethanol from tarpaulin sheets laid under nesting trees or other substrates on Narcondam Island between December 2019 and February 2020. PP and RN collected Wreathed Hornbill tissue samples in absolute alcohol from hornbill trophies preserved by local community members in Tirap, Changlang and Lower Dibang Valley districts of eastern Arunachal Pradesh. The hornbill trophies are displayed in local houses as part of their culture.

### DNA Extraction

Genomic DNA was extracted from the faecal samples using a Phenol:Chloroform:Isoamyl alcohol protocol developed for non-invasive samples of Bucerotidae (Hutton & Dingle, pers comm), modified from Coffroth et al. (1992). See supplementary materials for detailed description of faecal extraction protocol. The pellet was suspended in 30-60 µl of sterile Milli-Q, and Nanodrop readings were taken to measure DNA concentration.

DNA was kept at -20°C for long-term storage. Since most of the undiluted genomic DNA extracts were heavily pigmented, we additionally tested the use of genomic DNA purification kits (Monarch genomic DNA purification kit, New England Biolabs and NucleoSpin gDNA Clean-up kit, Macherey-Nagel) in the purification of DNA and in improving the efficiency of PCR amplification. DNA was extracted for 58 unique Narcondam Hornbill individuals using this procedure. DNA extractions from 19 unique Wreathed Hornbill tissue samples were done using Qiagen DNEasy Blood and Tissue kit following the manufacturer’s protocol, after which they were kept at -20°C for storage.

### PCR Amplification and Sequencing

The Narcondam Hornbill DNA extracts were amplified for Cytochrome B (Cytb; ∼500 bp), Cytochrome oxidase I (COI; ∼700 bp), NADH dehydrogenase subunit-2 (ND2; ∼1,400 bp) and non-protein coding D-loop region (D-loop; ∼500 bp), the latter for which a new primer pair was designed for this study. The D-loop marker, which is a faster evolving region, has been used in population genetics studies previously. The amplification was carried out for all faecal DNA extracts. However, the success rate varied among the samples. Sequences were obtained in at least one direction for 14 samples for Cytb and 28 for the D-loop region. COI and ND2 regions were amplified and sequenced successfully for 14 and 13 samples, respectively, with all four markers sequenced for 11 individuals. The PCR conditions and primer details are mentioned in SI Tables 1 and 2.

We selected 19 Wreathed Hornbill samples from eastern Arunachal Pradesh (Fig. 1), specifically Tirap, Changlang and Eastern Dibang Valley districts, for PCR amplification and sequencing of the same markers. The three districts were chosen in order to restrict sampling to a relatively small distribution range to minimise the potential biases caused by sampling across a wider geographic area, while also getting a comparable number of samples (13-14) to the successful Narcondam Hornbill sequences for the markers. They also had variable success with PCR amplification and sequencing. We could not sequence the D-loop region. The Cytb, COI and ND2 regions were successfully sequenced for 12, 16 and eight individuals, respectively.

### Assessing genetic diversity

The sequences were cleaned using SeqScanner2 and Chromas. The cleaned sequences were aligned using the MUSCLE (Edgar 2004) algorithm in MEGA 7 software (Kumar et al. 2016). The sequence statistics of these alignments were calculated in R 4.2.2 using functions from the packages ‘ape’ (Paradis et al. 2004), and ‘ips’ (Heibl et al. 2014). This included the detection of segregating sites, parsimony informative sites, nucleotide diversity (π, average pairwise difference) and haplotype diversity. Haplotypes were detected using the function *haplotype* from the ‘pegas’ package and exported as a nexus file. Since some sequences with missing data in indeterminate bases (N) were detected as unique haplotypes, redundant haplotypes were removed by manual inspection to keep unique haplotypes. Minimum-spanning haplotype networks were created using the PopART software (Leigh and Bryant 2015).

## RESULTS

The Narcondam Hornbill samples sequenced showed no detectable variation in the Cytb and COI regions. The Wreathed Hornbill samples exhibited three haplotypes in the Cytb region and seven in the COI region (Fig. 2). In the more variable ND2 locus, Narcondam samples had 10 variable sites and seven distinct haplotypes (Hd = 0.795), while Wreathed Hornbills, with a much lower sample size, had 13 segregating sites and seven haplotypes (Hd = 0.893) (Table 1). The nucleotide diversity for this region was 0.0069 for Wreathed Hornbills and 0.003 for Narcondam Hornbills. The D-loop region, only successfully sequenced for the Narcondam Hornbills, had four haplotypes.

**Figure 2.**
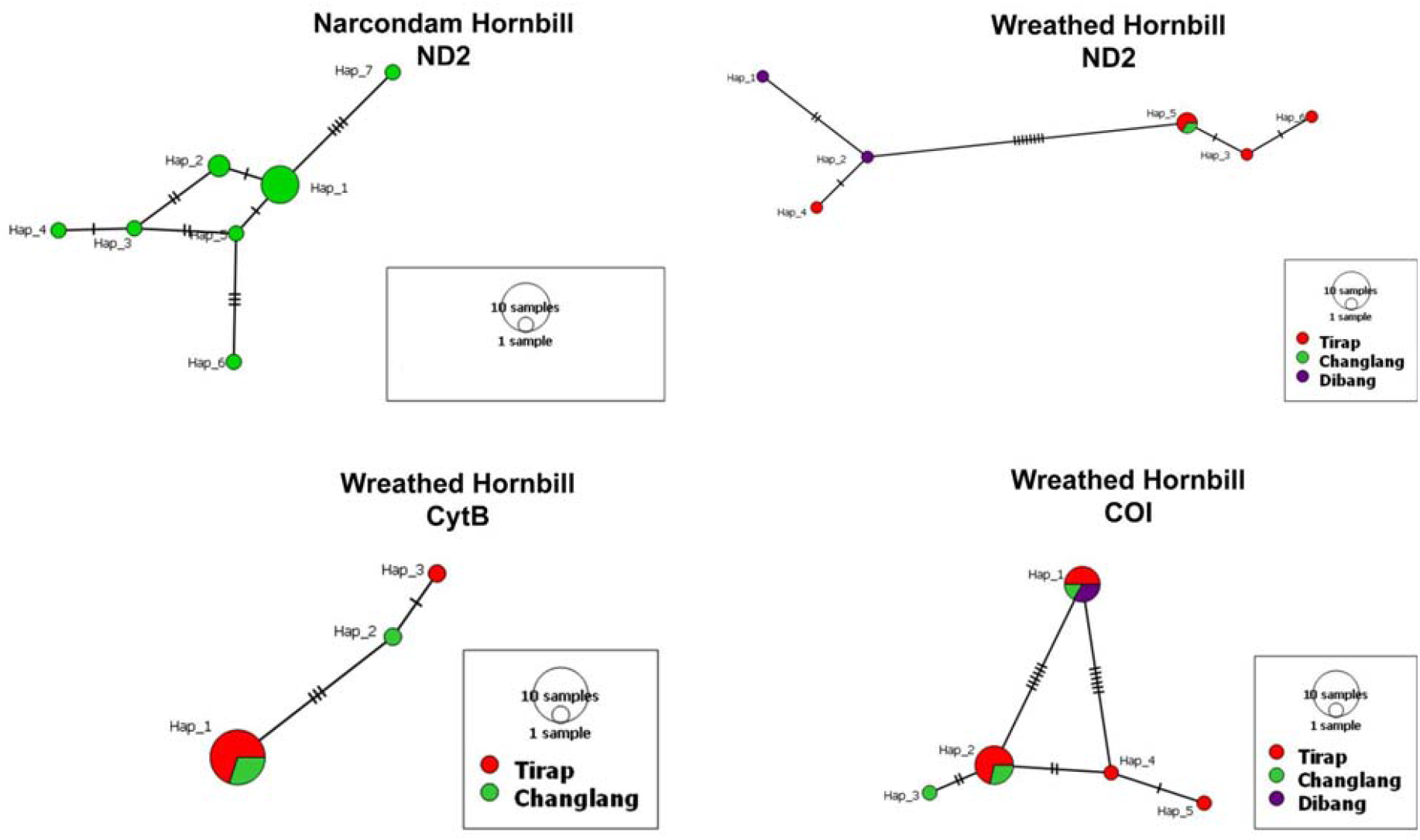
Minimum-spanning Haplotype networks of Narcondam and Wreathed Hornbills for markers where distinct haplotypes were identified.

**Table 1.** Genetic diversity indices for Narcondam Hornbill and Wreathed Hornbill samples that were successfully sequenced for four markers (Cytb, COI, ND2 and D-loop region).

|  | **Wreathed Hornbill** |  |  | **Narcondam Hornbill** |  |  |  |
| --- | --- | --- | --- | --- | --- | --- | --- |
|  | **Cytb** | **COI** | **ND2** | **Cytb** | **COI** | **ND2** | **D-loop** |
| **No. of sequences** | 12 | 16 | 8 | 14 | 14 | 13 | 28 |
| **Sequence length**<br>**in base pairs**<br>**(bp)** | 363 | 670 | 825 | 431 | 528 | 819 | 385 |
| **No. of**<br>**segregating sites** | 4 | 9 | 13 | 0 | 0 | 10 | 3 |
| **No. of parsimony**<br>**informative sites** | 3 | 7 | 9 | 0 | 0 | 4 | 0 |
| **Nucleotide**<br>**diversity** | 0.003 | 0.0054 | 0.0069 | 0 | 0 | 0.003 | 0.0006 |
| **No. of**<br>**haplotypes** | 3 | 5 | 6 | 1 | 1 | 7 | 4 |
| **Haplotype**<br>**diversity** | 0.318 | 0.700 | 0.893 | 0 | 0 | 0.795 | 0.206 |

## DISCUSSION

The island-endemic Narcondam Hornbill had very low genetic diversity compared to the more widespread Wreathed Hornbill. This could be because of three non-mutually exclusive reasons. First, it could be related to a relatively recent founder event (crown age 2.58 million years) that led to the origin of the species which now populate Narcondam Island (Gonzalez 2012). The colonisation of the Narcondam island may be associated with relatively few dispersal events, given the limited connectivity with mainland south-east Asia and evidence for limited overwater gene flow across a narrow channel in another insular hornbill (Sammler et al. 2012). More importantly, the long generation time of hornbills would mean that the Narcondam hornbill population likely remained in a bottleneck for a longer period of time negatively influencing the accumulation of genetic diversity since the founder event (Stuessy et al. 2014).

Second, the Narcondam hornbill is endemic to an island with a very small area (6.8 km^2^) as compared to its more widespread sister species (*Rhyticeros plicatus*). This can influence the population size that can be supported on the island and may lead to greater demographic fluctuations brought about by stochastic events. Genetic diversity is lower for species that have historically had smaller suitable habitats (Brüniche-Olsen et al. 2021), and a significant association between range size, population size, and genetic diversity has been recently demonstrated (Buffalo 2021).

Third, extensive hunting of Narcondam hornbill individuals was documented on the island in 1998, and the estimated population size of the Narcondam hornbill was around 300-400 birds (Sankaran et al. 2000; Yahya and Zarri 2002; Ramachandran 2022). In contrast, the most recent estimates, when hunting pressures are absent, suggest a population size of around 1,000 birds (Naniwadekar et al. 2021). This indicates significant declines in population size due to hunting in the past that could have resulted in a recent decline in genetic diversity.

The nucleotide diversity observed in the mitochondrial D-loop region for 28 Narcondam hornbill samples (0.0006) is lower than values reported for island populations of *Penelopides panini* (0.008-0.015), *P. manillae* (0.007, 0.037), *Rhabdotorrhinus waldeni* (0.005, 0.011) and *R. leucocephalus* (0.038) within the Philippine archipelago (Sammler et al. 2012). These species belong to the same insular clade of Asian hornbills as the Narcondam Hornbill and, therefore, make a reasonable point of comparison. Among the Philippine hornbills, *P. panini* and *R. waldeni* also show a loss of haplotypes as compared to their more abundant relatives, which has been related to a recent decline in population size due to hunting and habitat destruction. Unfortunately, we were unable to extract D-loop information for the Wreathed Hornbill.

We successfully obtained mtDNA sequences from Narcondam Hornbill faecal samples. The limited success of amplification across samples may be related to the presence of uric acid and plant matter and their secondary metabolites, which hinder the extraction of genomic DNA (Wilson 1997; Munch et al. 2019). The efficiency of amplification improved marginally when an additional step of column-based purification was added to the extraction of genomic DNA. However, variability in PCR success across samples persisted. Given that most wild, forest-dwelling hornbill species are challenging to capture, faecal samples can be a useful source of hornbill DNA.

Theory, field and lab observations show that low genetic diversity is associated with a decline in population viability and adaptive potential, which could lead to extinction (DeWoody et al. 2021; Kardos et al. 2021). For most organisms, we lack the ability to specifically measure and conserve functional genetic diversity that underlies phenotype and, therefore, individual fitness. In this case, it is important to monitor and conserve genome-wide variation and the processes that facilitate it to conserve Narcondam Hornbill in the long term.

## ACKNOWLEDGEMENTS

We thank Wildlife Conservation Trust, Mr. Uday Kumar, M. M. Muthiah Research Foundation, Mr. Rohit and Deepa Sobti, and Mr. Aravind Datar for providing funding support. We thank Arunachal and Andaman & Nicobar Forest and Police Departments, Indian Coast Guard, for giving us permission to conduct the study. We thank Mr. D. M. Shukla, Commandant A. K. Bhama, Captain Kundan Singh, Mr. Dependra Pathak, Ms. Usha Rangnani, Mr. A. K. Paul, Mr. Soundra Pandian, Mr. Bhatt, Mr. Damodhar A. T., Mr. Harshraj Wathore, Mr. Tana Tapi, Mr. Kime Rambiya, Mr. Diwang Lowang, Mr. Mayur Wariya, Mr. Chandan Ri for support during fieldwork. We thank Payal Dash for assistance with lab work and thank the DNA sequencing facility at CSIR-Centre for Cellular and Molecular Biology. We are grateful to the then Dean and Director WII, Dr. G. S. Rawat, for supporting the project. We would like to thank Mr. Japang Pansa and Mr. Santosh Barua for logistical support during the study. We thank Caroline Dingle and Chloe Hutton for sharing DNA extraction protocols. We thank Elrika D’Souza, Evan Nazareth, Rachana Rao and Rohan Arthur for providing support in Port Blair. We thank Prasenjeet Yadav, Suri Venkatachalam, Shashank Dalvi, and Anand Osuri for valuable discussions.

## SUPPLEMENTARY MATERIAL

### Extraction Protocol for faecal samples

About 200 mg of faecal matter was spun down and the ethanol removed, followed by washing with 300 µl of 2X CTAB digestion buffer (1.4 M NaCl, 20 mM EDTA, 100 mM Tris-HCL, 2% w/v CTAB, immediately before digestion addition of 0.2% v/v β-mercaptoethanol and 2% w/v polyvinylpyrrolidone to the buffer). Digestion was carried out in 600 µl of 2X CTAB buffer and 20 µl of 20 mg/ml proteinase K, where samples were incubated at 65°C for 24-48 hrs with occasional vortexing. To the digested samples, 600 µl of 24:1 Chloroform:Isoamyl alcohol was added and vortexed, followed by centrifugation. To the aqueous layer, 600 µl of 25:24:1 Phenol:Chloroform:Isoamyl alcohol was added and vortexed followed by centrifugation.

Chloroform:Isoamyl alcohol extraction was repeated with the aqueous layer to remove any remnant contamination with phenol. Finally, DNA was precipitated by the addition of 1 ml of ice-cold 95% ethanol to the aqueous layer and incubated at -80°C for 30 minutes. After centrifugation, the DNA pellet was washed twice with 500 µl of ice-cold 70% ethanol by inversion. The pellet was allowed to air dry after removing any remnant 70% ethanol with a pipette. The pellet was suspended in 30-60 µl of sterile Milli-Q, and Nanodrop readings were taken to measure DNA concentration.

#### Primers Used

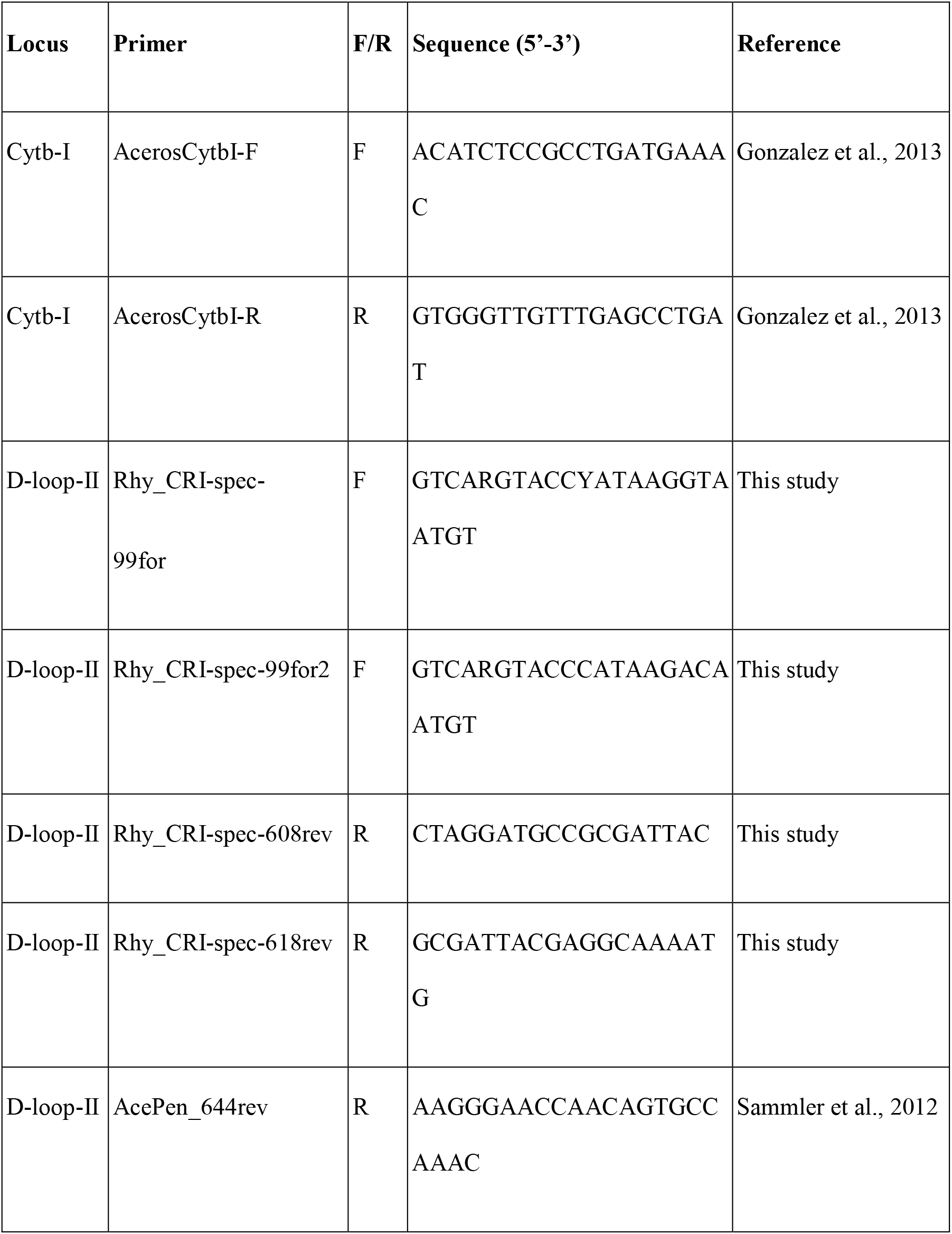
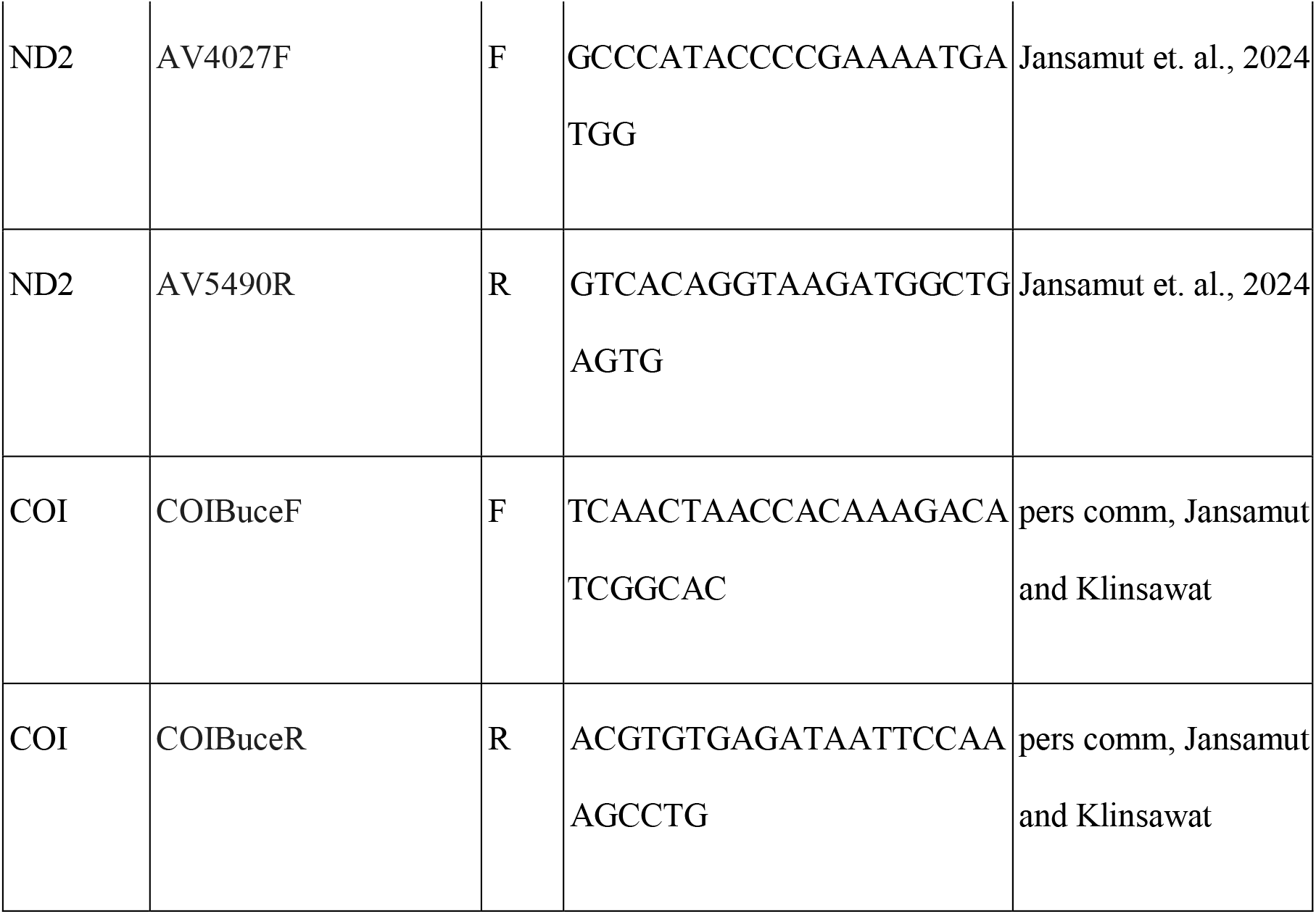

#### PCR Conditions

| Primer Pair |  | Locu<br>s | Prod<br>uct<br>Size | Denatu<br>ration | Anne<br>aling | Exten<br>sion | Final<br>Extens<br>ion | Num<br>ber<br>of<br>Cycle<br>s | Prim<br>er<br>Conc | Enhan<br>cers<br>(Conc<br>in<br>reactio<br>n<br>mixtur<br>e) |
| --- | --- | --- | --- | --- | --- | --- | --- | --- | --- | --- |
| Forwa<br>rd | Rever<br>se |  |  |  |  |  |  |  |  |  |
| Acero<br>sCytbI<br>-F | Acero<br>sCytbI<br>-R | Cytb | ~500 | 94□,<br>30s | 53□,<br>30s | 72□,<br>45s | 72□,<br>5m | 30 | 30% | BSA<br>(20%)<br>MgCl<sub>2</sub><br>(1-2mM)<br>DMSO<br>(6-8%) |
| Rhy_<br>CRI-spec-99for2 | Rhy_<br>CRI-spec-618re<br>v | DLoo<br>p | 519 | 94□,<br>30s | 49□,<br>30s | 72□,3<br>0s | 72□,<br>5m | 35 | 10% | BSA<br>(1.3%) |
| Rhy_<br>CRI-spec-99for2 | AcePe<br>n_deg<br>—<br>644re<br>v | DLoop | 540 | 94□,<br>30s | 49-50.5□<br>,<br>45s | 72□,3<br>0s | 72□,<br>5m | 40-50s | 20% | BSA<br>(1.3-2.3%) |
| COIB<br>uceF | COIB<br>uceR | COI | ~700 | 94□,<br>30s | 53□,<br>30s | 72□,<br>45s | 72□,<br>5m | 35 | 30% | BSA<br>(20%)<br>MgCl2<br>(1-2mM)<br>DMSO<br>(6-8%) |
| AV40<br>27F | AV54<br>90R | ND2 | 1400 | 94□,<br>30s | 53□,<br>30s | 72□,<br>45s | 72□,<br>5m | 35 | 30% | BSA<br>(20%)<br>MgCl2<br>(1-2mM)<br>DMSO<br>(6-8%) |

